# Scalable Extraction of Information on Protein-Protein Interactions using Topological Data Analysis

**DOI:** 10.64898/2026.08.06.743405

**Authors:** Angan Mukherjee, ByungUk Park, Adam Malmstrom, Jessi Cisewski-Kehe, Reid C. Van Lehn, Victor M. Zavala

## Abstract

Protein-protein interactions (PPIs) govern a wide range of cellular functions. The ability to predict PPI interfaces from protein molecular surfaces is important for understanding protein function and enabling therapeutic discovery. While recent advances in structure-based learning, particularly molecular-surface geometric deep learning frameworks, have demonstrated that protein surfaces encode rich geometric and physicochemical information, such approaches often remain computationally intensive and data-hungry. Alternatively, topological data analysis (TDA) has emerged as a mathematically rigorous framework for extracting robust, multiscale shape information from complex data. In this work, we introduce a scalable TDA framework for extracting information on PPIs directly from localized protein surface patches. Our approach leverages multiscale topological descriptors, evaluated from patch-wise point cloud representations of protein mesh surfaces, combined with supervised machine learning models for interface prediction. On a full dataset of 3,362 proteins, the proposed approach substantially reduced computational cost relative to an established geometric deep learning method, MaSIF-site, decreasing preprocessing time from approximately 27 s/protein to 5-8 s/protein and total training time from approximately 6 h to 1-1.3 h. Importantly, this computational reduction is achieved while maintaining mean test area under the receiver operating characteristic curve (AUC) values of 0.76 and 0.77 for patch radii of 9 Å and 12 Å, respectively, thus approaching the MaSIF-site test AUC of 0.84. Our results suggest that topology offers a scalable and computationally efficient approach for high-throughput extraction of information from complex biomolecular interfaces.

## 1 Introduction

Protein-protein interactions (PPIs) are the primary drivers for cellular organization and function, mediating processes ranging from metabolic regulation to immune responses [1, 2]. Accurate prediction of PPI interfaces is critical for identifying *druggable* surface regions and guiding the design of molecules that modulate protein binding [3]. To address this challenge, specialized PPI modeling frameworks have been developed to enable rapid and scalable identification of interaction-prone or druggable regions on target proteins [4, 5, 6, 7, 8]. These methods typically leverage graph-based descriptions of protein structure to learn patterns associated with protein-protein recognition.

At the same time, the broader revolution in protein-structure prediction has reshaped the context in which interface prediction is performed. Co-folding approaches based on AlphaFold-style architectures [9], including AlphaFold2-derived multimer methods [10] and more recent AlphaFold3 formulations [11], have achieved remarkable progress in predicting the structures of interacting biomolecular complexes. Advances in protein foundation models have further enabled the *de novo* design of high-affinity binders against diverse biomolecular targets [3, 11, 12, 13, 14, 15, 16].

Despite these advances, co-folding methods do not eliminate the need for dedicated interface representations or fast, scalable approaches for identifying interaction-prone regions on individual protein structures. Recent studies have noted that co-folding-based strategies can remain limited by specificity, dependence on successful complex formation, system type, or computational cost relative to direct interface prediction from known structures [9, 5, 17]. Thus, methods that directly characterize interaction-prone regions from a single protein structure remain valuable, particularly when scalability, interpretability, and integration into drug-discovery workflows are important [5].

Geometric deep learning has emerged as a powerful framework for processing three-dimensional protein structures and identifying interaction-prone regions. For example, ScanNet [7] and PeSTo [5] have shown that structure-based learning can successfully predict binding interfaces by operating on local three-dimensional neighborhoods of atoms or residues, often improving interpretability, interface generality, or computational efficiency. A related surface-centered framework, molecular surface interaction fingerprinting (MaSIF) [4], represents biomolecular surfaces as collections of local patches described by numerical geometric and chemical descriptors. The MaSIF framework includes task-specific models, including MaSIF-site [4], which predicts protein-protein interaction sites from local molecular surface patches. MaSIF has demonstrated broad applicability across diverse molecular interaction contexts, including protein-protein, protein-small molecule, and protein-membrane interactions [4, 18, 19]. It has also enabled the *de novo* design of peptides that bind specific protein surfaces [20, 21]. More recently, MaSIF-based approaches have been used to design peptides targeting novel binding sites, or ‘neosurfaces’, that emerge upon protein-ligand complex formation, thereby expanding the targetable space for drug discovery [22, 23]. In addition, protein surface representations derived from MaSIF-like frameworks provide useful tools for fragment-based drug discovery and for mapping druggable sites across the human surfaceome to support the design of novel binders [24, 25]. Together, these studies highlight the generalizability of surface fingerprint descriptors for predicting molecular interactions and guiding *de novo* binder design.

However, surface-centered deep learning methods still present some limitations that motivate alternative data representations. First, many such methods remain relatively data hungry, with learned convolutional or point-based feature extractors requiring substantial training data and computational effort to capture the spatial invariants needed for robust generalization. Second, surface parameterization, meshing, and feature mapping can introduce nontrivial computational preprocessing over-head and sensitivity to representation details [5]. Third, although these approaches often achieve strong predictive accuracy, their learned embeddings are not always compact or directly interpretable in terms of physically meaningful structural organization. These limitations become increasingly important when one seeks high-throughput screening across large datasets, repeated analysis over trajectories or conformational ensembles, as well as integration into workflows where computational efficiency and representation simplicity matter [26]. Such concerns have been explicitly recognized even in recent state-of-the-art methods, which continue to emphasize speed, scalability, and robustness as active design objectives [18].

Topological data analysis (TDA), specifically persistent homology (PH), has emerged as a powerful alternative for characterizing the shape and structure of complex datasets in chemical engineering and materials science [27, 28, 29]. TDA provides a framework for converting complex molecular data into robust multiscale descriptors that summarize shape, connectivity, loops, and related structural organization [30]. A key strength of TDA is its ability to reduce high-dimensional data into low-dimensional, interpretable descriptors such as Euler characteristics (ECs), Betti numbers [30, 31], and persistence landscape functions (PL) [32, 33]. The interpretability of these descriptors arises from their direct topological meaning: EC and Betti functions summarize how connected components, loops, and void-like structures emerge and disappear across filtration scales. In the context of localized protein surface patches, these quantities provide compact summaries of multiscale surface organization, roughness, and cavity-like geometric structure. Topological descriptors complement traditional geometric descriptors by explicitly characterizing the evolution of connected components, loops, and voids across a *filtration* of spatial scales. In particular, PH representations provide a systematic approach to summarize multiscale structural organization and admit formal stability guarantees under perturbations of input data [34, 35].

TDA has also gained popularity in biomolecular modeling as an efficient framework for extracting multiscale structural information from high-dimensional molecular data based on chemical heterogeneity [36, 37, 38, 34, 39, 40]. For example, researchers have implemented an integrated framework that combines element-specific PH with machine learning (ML) approaches to predict proteinligand binding affinity [41]. Element-specific PH retains the geometric information encoded by the filtration while incorporating chemical identity by constructing TDA descriptors for selected atom types or atom-type combinations, thereby yielding a more chemically informed representation of biomolecular interactions [42]. More recently, weighted PH formulations have incorporated physical, chemical, and biological information directly into topological constructions, enabling the joint representation of geometry and molecular heterogeneity in a unified framework [43]. These studies demonstrate that TDA can provide compact and informative descriptors for biomolecular systems; however, its use has remained focused primarily on structural characterization and *global* molecular property prediction, rather than localized prediction of PPI from molecular surface patches. Particularly, its use has not been systematically developed for localized, surface-centric prediction of protein-protein interfaces.

In this work, we address this gap by introducing a scalable TDA framework for PPI interface prediction directly from localized protein surface patches. Starting from the atomic coordinates of the protein, we generate a discretized molecular dot surface and construct a triangulated mesh representation, with each surface point assigned either an interface and non-interface label. Each surfacemesh vertex is used as a patch center, and the associated local point cloud consists of all surface vertices located within a fixed Euclidean radius of that center. Topological summaries are then extracted from each overlapping local patch using a moving-patch approach. As illustrated in Figure 1, each patch is represented as a 3D point cloud from which simplicial complexes are constructed to compute multiscale topological descriptors, such as EC and Betti functions. We further explore multiple representations of localized protein surface meshes through different simplicial complexes (e.g., alpha, rips, and cubical complexes), together with different topological summaries derived from PH (e.g., EC and Betti numbers) and PL. These descriptors are then augmented with local chemical features, such as electrostatics, hydrogen-bond potential, and hydrophobicity, to preserve both geometric/topological and physicochemical information relevant to intermolecular recognition. The combined descriptors are fed to a supervised learning model, represented by a multilayered feedforward neural network (NN), for prediction of PPI sites. In this way, the proposed framework offers a compact and scalable alternative to data-hungry geometric deep learning pipelines, typically associated with significantly higher computational expense, while *mostly* preserving the localized, surface-based structural representations that have proven successful in MaSIF- and dMaSIF-type approaches [4, 18, 20, 22, 19].

**Figure 1.**
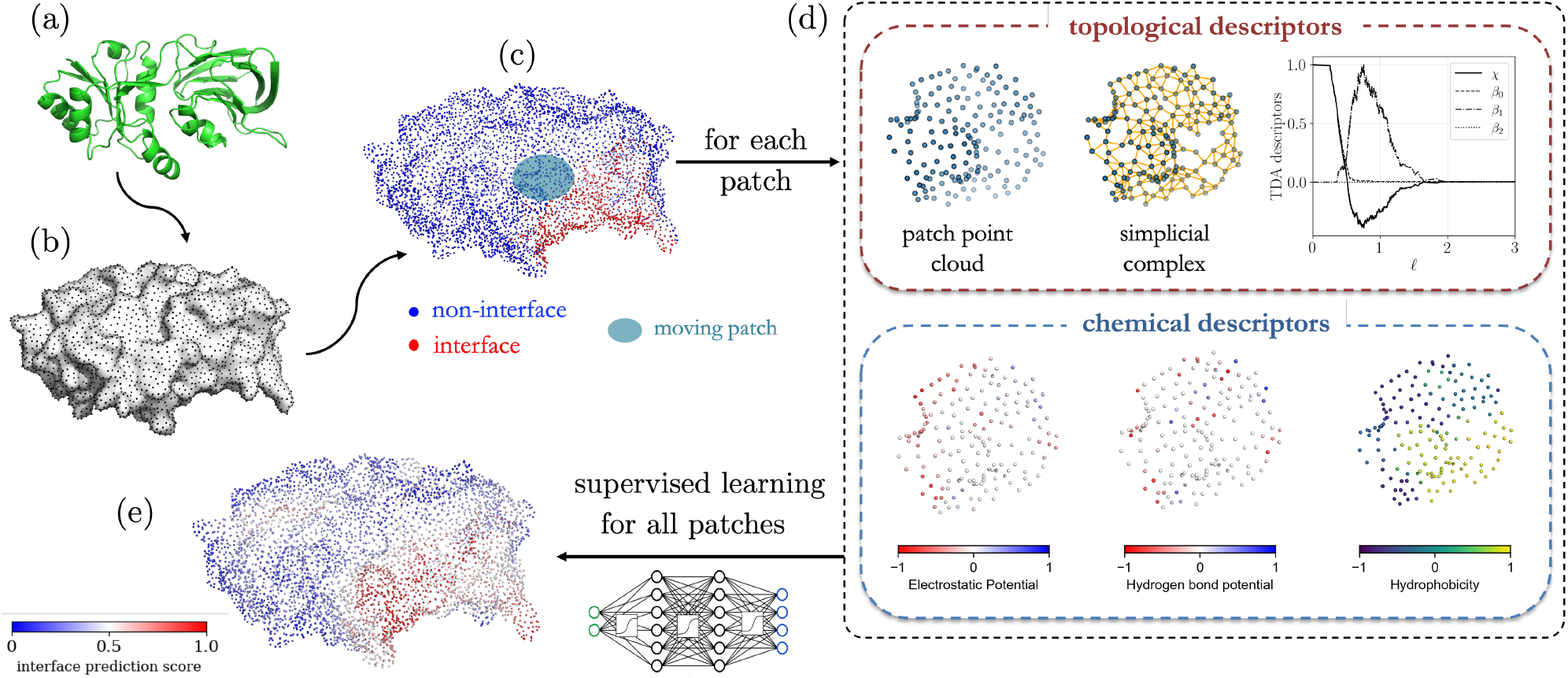
Proposed TDA framework for PPI prediction. (a) A 3D protein structure obtained from atomic coordinates. (b) Protein atomic coordinates are used to generate a molecular surface representation. (c) For labeled benchmark structures, the surface-mesh vertices are annotated as interface or non-interface points, and overlapping local surface patches of fixed radius are extracted around each vertex, with each patch represented as a point cloud. (d) TDA descriptors are obtained by constructing a simplicial complex for each patch and computing topological summaries such as EC (*χ*) and Betti numbers (*β*_0_, *β*_1_, *β*_2_). The topological descriptors are augmented with chemical descriptors, such as electrostatics, hydrogen-bond potential, and hydrophobicity. (e) The combined set of topological and chemical descriptors are used in a supervised learning model to predict interface and non-interface scores, which are projected back onto the molecular surface to identify PPI sites.

The paper is organized as follows. Section 2 presents the mathematical and computational foundations of the proposed framework. Section 3 evaluates the proposed methodology on benchmark PPI datasets and compares its predictive performance against representative geometric deep learning-based fingerprinting approaches (such as MaSIF). Section 4 summarizes the key findings from this work, discusses current limitations, and outlines directions for future research.

## 2 Computational Workflow

We introduce the detailed computational elements of the proposed workflow for PPI interface prediction. The overall methodology can be categorized into the following stages: (a) construction of localized molecular-surface patches, (b) extraction of TDA descriptors using a moving-patch point cloud representation, (c) dimensionality reduction of the combined descriptor space involving topological and radially averaged chemical features using principal component analysis (PCA), and (d) training of a supervised learning model (NN) on the augmented feature (input) set for predicting PPI interfaces. Detailed descriptions of the dataset under consideration, computational preprocessing for dot surface generation, chemical feature computation, and interface label assignment are included in the *Supporting Information* document.

### 2.1 Moving-Patch Approach

Let 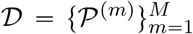 denote a dataset of *M* ∈ ℤ_+_ proteins obtained from the RCSB protein data bank (PDB) [44], where each protein *P* ^(*m*)^ is represented by the atomic coordinates of its heavy atoms. The corresponding dot surface for each protein is generated from the atomic coordinates using `MSMS` [45] for the moving-patch construction. Additional details on the computation of the discretized molecular dot surfaces are provided in Section S.2.1 of the *Supporting Information* document. In this work, *M =* 3362. Additional information on the PPI dataset considered in this work can be found in Section S.1 of *Supporting Information*. For each protein *P*^(*m*)^, let

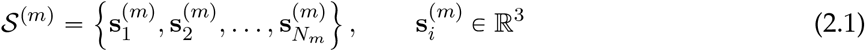

denote the set of discretized molecular-surface points, where *N*_*m*_ ∈ ℤ_+_ is the number of surface points for protein *m*. A binary label 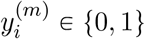 is associated with each surface point 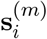, where 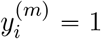 indicates that the surface point belongs to an interface region and 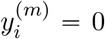 indicates that it belongs to a non-interface region, as discussed in Section S.2.3 of *Supporting Information*.

To analyze the local structural neighborhood of each surface point, we construct overlapping surface patches using a spherical moving-patch approach. Specifically, for patch radius *R*_p_ > 0, the patch centered at 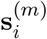 is defined as:

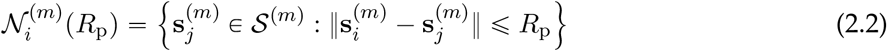

where ∥ · ∥ denotes the Euclidean distance in ℝ^3^. Thus, each surface point 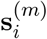 serves as the center of a local patch, and the corresponding patch label is assigned as the label 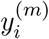 of the center point.

Additionally, let

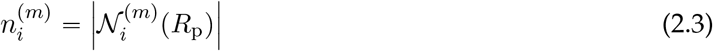

denote the number of points in the *i*-th patch of the *m*-th protein. Therefore, each local patch is represented as a finite point cloud, defined as:

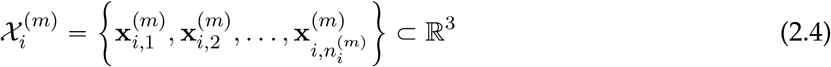

where 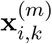 denotes the Cartesian coordinate vector of the *k*-th surface point in that patch, for 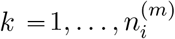. In this work, the TDA descriptors are computed directly from these localized point cloud representations of the molecular surface.

### 2.2 Extraction of Topological Information

For notational simplicity, we consider a generic local patch point cloud, defined as:

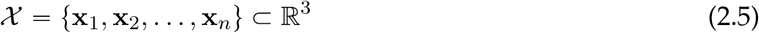

where *n* ℝ ℤ_+_ denotes the number of surface points in the patch and **x**_*k*_ ∈ ℝ^3^ denotes the Cartesian coordinate vector of the *k*-th point in the patch. We implement TDA to extract multiscale descriptors that summarize the geometric and topological structure of *X*.

#### 2.2.1 Topological Representations of Surface Geometry

In this work, we consider multiple topological representations of the patch point cloud *X* through simplicial-complex constructions [46], such as Vietoris-Rips (VR) and alpha complexes, as well as cubical complexes [47]. All topological descriptors are computed using the default implementations available in the `GUDHI` library [48] in Python.

##### Vietoris-Rips (VR) complex

The VR complex provides a way to convert the patch point cloud into a multiscale combinatorial representation by connecting points that are sufficiently close to one another. As the filtration threshold *ℓ* increases, additional connections and higher-dimensional structures are added to the representation.

To describe these structures, we introduce the notion of a simplex. A simplex is a basic geometric object defined by a set of vertices: a 0-simplex is a vertex, a 1-simplex is an edge connecting two vertices, a 2-simplex is a filled triangular face connecting three vertices, and a 3-simplex is a tetrahedron connecting four vertices. Additionally, a simplicial complex is a collection of simplices that also contains all lower-dimensional faces of each included simplex. For example, if a 2-simplex is included, then its three edges and three vertices are also included.

To define the VR complex, a distance between vertices is needed. Let

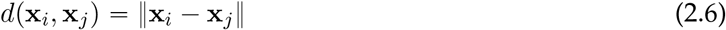

denote the Euclidean distance between two patch points **x**_*i*_, **x**_*j*_ ∈ *χ*. For a filtration value *ℓ* ∈ ℝ_+_, the VR (rips) complex [49] of *X* is defined as:

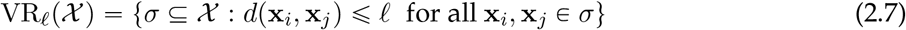

where *σ* denotes a simplex whose vertices are points from *X*. Thus, a simplex of any dimension is included in VR_*ℓ*_(*χ*) whenever all pairwise distances among its vertices do not exceed the threshold *ℓ*. For example, two points within distance *ℓ* define a 1-simplex together with its two 0-simplex faces, while three points that are all pairwise within distance *ℓ* define a 2-simplex together with its three edges and three vertices. The family tVR_*ℓ*_(*χ*)u_*ℓ*ě0_, therefore, defines a filtration as:

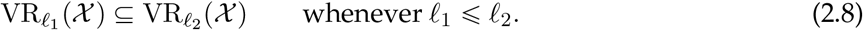

This construction is illustrated in the top row of Figure 2: as *ℓ* increases, balls centered at the point cloud locations begin to overlap, edges are introduced between nearby points, and higher dimensional simplices are added when groups of points are mutually within the distance threshold.

**Figure 2.**
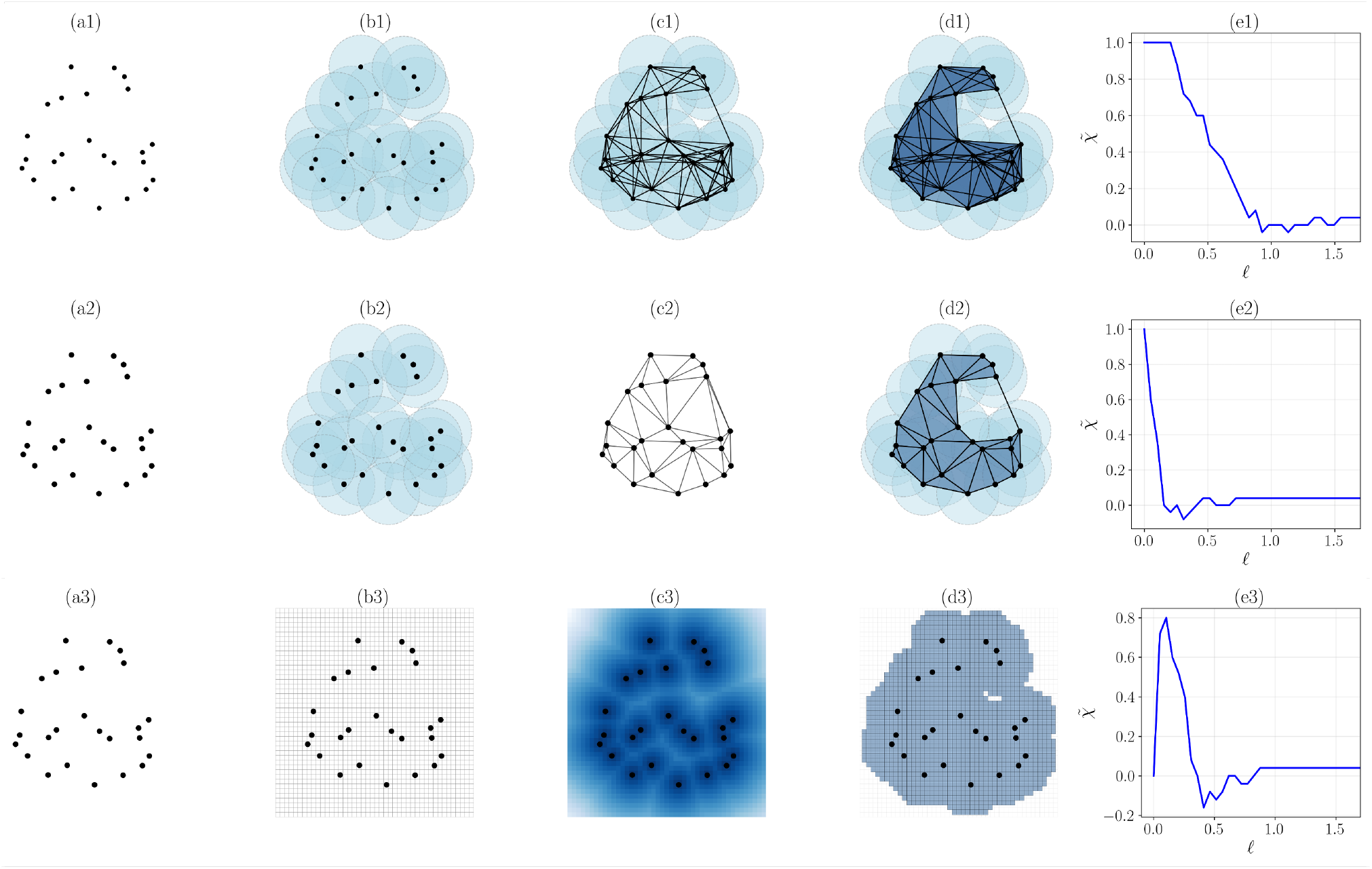
Illustration of the computation of EC functions from the rips (top row), alpha (middle row), and cubical (bottom row) complex representations of a sample 3D point cloud. (a1), (a2), and (a3) show the same representative point cloud. For the rips complex, (b1) shows balls of radius *ℓ*/2 centered at the sample points, (c1) shows the corresponding 1-skeleton formed by edges between points whose pairwise Euclidean distance is at most *ℓ*, (d1) shows the resulting rips complex with higher-order simplices (filled triangular faces), and (e1) shows the resulting EC function normalized by the number p*n*q of points in the 3D point cloud 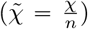, computed from the filtration operation. For the alpha complex, (b2) shows the union of balls used in the geometric construction, (c2) shows the Delaunay triangulation of the point cloud, (d2) shows the corresponding alpha complex obtained as a sub-complex of the Delaunay triangulation, and (e2) shows the normalized EC function. For the cubical complex, (b3) shows the regular cubical grid, (c3) shows the grid-based scalar-field representation used to define the cubical filtration, (d3) shows the associated sublevel-set cubical complex, and (e3) shows the EC function normalized by the number of points. The representative filtration thresholds used to generate the respective complexes are *ℓ* = 1.5 (rips), *ℓ* = 0.5625 (alpha), and *ℓ* = 0.75 (cubical), respectively. These representative filtration values are selected only to provide visually interpretable examples; their numerical values are not directly comparable across the three filtrations because the corresponding filtration parameters have different constructions and units. Although shown here for a representative 3D point cloud for visualization, the same approach is applied in this work to localized protein surface patches for computation of topological descriptors.

##### Alpha complex

Let Del(*χ*) denote the Delaunay triangulation [50, 51] of *X*. For a filtration value *ℓ* ∈ ℝ_+_, the alpha complex A_*ℓ*_(*χ*) is the sub-complex of Del(*χ*) containing simplices whose corresponding empty circumsphere radius is bounded by 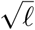. Equivalently, A_*ℓ*_(*χ*) may be viewed as the nerve of the intersection of Voronoi cells [52, 53] with balls of radius 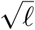 centered at the points in *χ*. The family {A_*ℓ*_(*χ*)}_*ℓ*ě0_ thus forms a filtration operation, defined by:

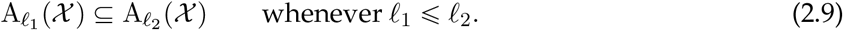

Compared to rips complexes, alpha complexes exploit the ambient Euclidean geometry more directly and often provide a sparser representation of the same point cloud [54]. The corresponding geometric construction is shown in the middle row of Figure 2, where the alpha complex is obtained as a filtration of the Delaunay triangulation and therefore retains only geometrically admissible simplices associated with the underlying point cloud arrangement.

##### Cubical complex

To construct a cubical representation of the point cloud, the localized patch geometry is embedded into a regular three-dimensional grid Ω ⊂ ℝ^3^ and a scalar field *g* : Ω → ℝ is defined on the grid. For a filtration threshold *ℓ* ∈ ℝ_+_, the associated sublevel-set cubical complex [47, 55] is defined as:

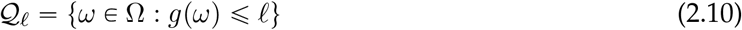

together with the collected set of vertices, edges, squares, and cubes on the grid. These elements are the cubical analogues of 0-, 1-, 2-, and 3-simplices, respectively, with squares and cubes replacing triangular faces and tetrahedra as the corresponding two- and three-dimensional building blocks. In this work, the scalar field is defined as the distance transform of the point cloud, i.e., 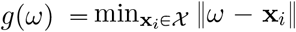 for each grid location *ω* ∈ Ω. Thus, *Q*_*ℓ*_ contains the grid cells whose distance to the nearest point in the patch is no greater than *ℓ*. As *ℓ* increases, the family {*Q*_*ℓ*_}_*ℓ*_ forms a nested sequence of cubical complexes and hence defines a cubical filtration. This grid-based construction is illustrated in the bottom row of Figure 2, where the point cloud is first embedded into a regular grid, converted into a distance-based scalar field, and then represented through a nested sequence of sublevel-set cubical complexes as *ℓ* increases.

#### 2.2.2 Topological Summaries

Let {*K*_*ℓ*_}_*ℓ* ∈ ℒ_ denote a generic filtration associated with a patch, where ℒ denotes the filtration domain. Depending on the specific representation as discussed in Section 2.2.1, *K*_*ℓ*_ may correspond to a rips complex, an alpha complex, or a cubical complex. For each homological dimension *q* ∈ ℤ_+ ∪ {0}_, the *q*-th Betti number at filtration value *ℓ* is denoted by:

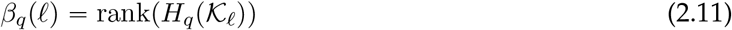

where *H*_*q*_ (*K*_*ℓ*_) is the *q*-th homology group of *K*_*ℓ*_. In particular, *β*_0_ (*ℓ*) refers to the number of connected components, *β*_1_p*ℓ*q denotes the number of one-dimensional holes or loops, and *β*_2_ (*ℓ*) signifies the number of enclosed voids within the 3D point cloud. Therefore, the EC [27, 31] at filtration value *ℓ* is defined as:

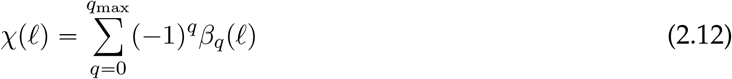

where *q*_max_ is the highest homological dimension retained in the computation. For the 3D patch point cloud considered here, we evaluate the EC as:

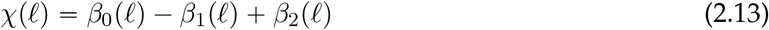

The construction of the three topological representations introduced in Section 2.2.1, along with the corresponding EC functions defined in Equation (2.12), is illustrated in Figure 2 for a representative sample 3D point cloud. Specifically, for a discretized filtration domain *ℓ*_1_ < *ℓ*_2_ < ⃛ < *ℓ*_*L*_, the EC and Betti functions for a given patch point cloud are represented by:

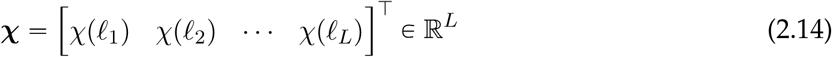

and

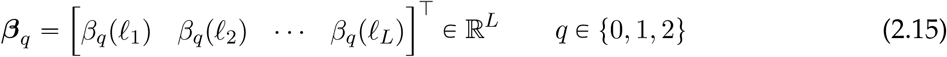

In addition to EC and Betti functions, we also consider PL [33, 56] as functional topological summaries of the 3D patch point cloud. For a given homological dimension *q*, let 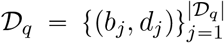 denote the corresponding persistence diagram, where *b*_*j*_ and *d*_*j*_ are the birth and death filtration values of the *j*-th topological feature. Each persistence pair defines a tent function:

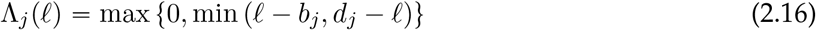

The *k*-th persistence landscape level is then defined point-wise as the *k*-th largest value among these tent functions:

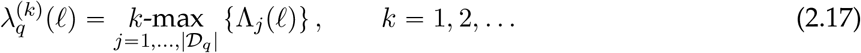

where *k* ∈ ℤ_+_ denotes the landscape level. Therefore, for each fixed homological dimension *q*, the PL provides a hierarchy of piecewise-linear functions over the filtration domain ℒ, with lower values of *k* capturing the most prominent topological features and higher values of *k* capturing progressively less dominant features.

For computational purposes, these landscape functions are sampled on the same discretized filtration grid *ℓ*_1_ < *ℓ*_2_ < ⃛ < *ℓ*_*L*_. Specifically, the discretized *k*-th landscape in homological dimension *q* is defined as:

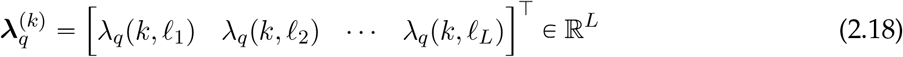

In practice, only the first few nonzero landscape levels are retained to obtain a finite-dimensional topological descriptor suitable for downstream learning. Figure 3 illustrates the construction of a PL from a representative sample 3D point cloud. Specifically, the figure shows how the topological descriptors identified across different homological dimensions (*q* = 0, 1) are transformed into piecewise-linear landscape functions (*λ*_*q*_) over the filtration domain. Accordingly, the proposed framework is not restricted to a single topological summary; rather, we investigate multiple summaries of localized protein surface geometry for optimal predictive capabilities.

**Figure 3.**
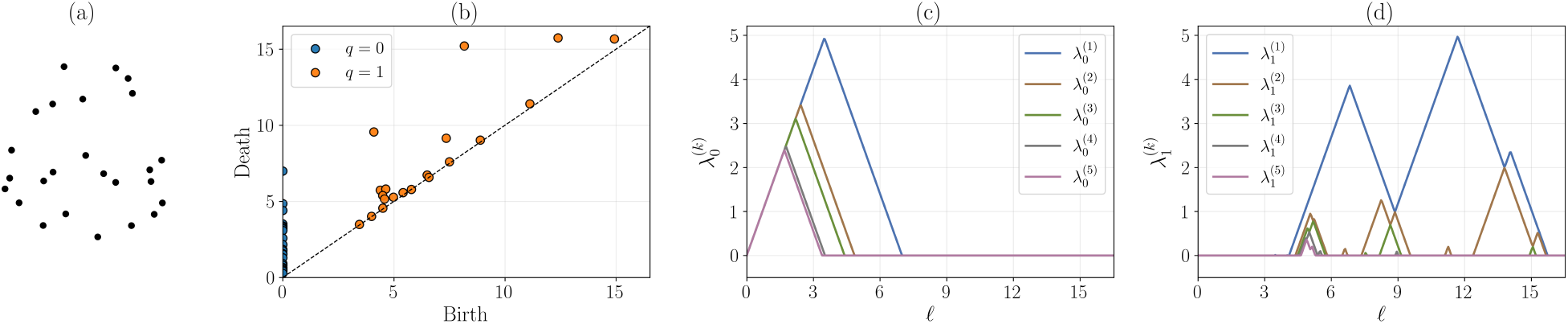
Illustration of persistence landscapes (PL) for (a) a representative sample 3D point cloud. (b) Persistence diagram obtained from the filtration of the point cloud, showing topological features in homological dimensions *q* = 0 and *q* = 1, where each point corresponds to the birth and death filtration values of a feature. The single *H*_0_ feature with infinite death time, corresponding to the connected component that persists throughout the filtration, is omitted for visualization and for PL computation. (c) PL functions 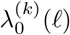 for homological dimension *q* = 0, showing the first five landscape levels as piecewise-linear summaries over the filtration domain. (d) PL functions 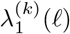 for homological dimension *q* = 1, also shown for the first five landscape levels. Lower landscape levels capture the most prominent topological features, while higher levels encode progressively less dominant features.

### 2.3 Dimensionality Reduction of Topological and Chemical Descriptors

The next step in the proposed workflow is to reduce the dimensionality of the combined patch-wise descriptor space involving topological and chemical features in order to improve computational tractability using PCA [57, 58]. Let 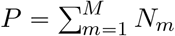, *P* ∈∈ ℤ_+_ denote the total number of surface patches across all proteins constructed from atomic coordinates obtained from the PDB dataset [44]. For each patch *p* ∈ {1, …, *P*}, let *χ*^(*p*)^ ∈ ℝ^*L*^ denote the discretized EC function defined in Equation (2.14). In addition, let *f*_elec_, *f*_hb_, *f*_hphob_ denote the electrostatic, hydrogen-bond, and hydrophobicity features defined on the molecular surface, respectively. Additional details on the calculation of chemical surface features are included in Section S.2.2 of *Supporting Information*. For each patch center, these chemical descriptors are averaged over nested radial neighborhoods, where each neighborhood contains all surface points within a prescribed radius from the center. Because these neighborhoods are cumulative rather than shell-wise, surface points closer to the patch center are included in multiple radial averages. For example, in this work, cumulative averages are computed over nested radial neighborhoods with radii of 0, 3, 6, and 9 Å for the 9 Å patch-radius case. The resulting averaged descriptors are concatenated to obtain a compact summary of the local physicochemical environment.

Let 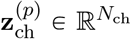 denote the resulting vector of radially averaged chemical descriptors for the *p*-th patch, where *N*_ch_ ∈ ℤ_+_ is the total number of available chemical features. Therefore, the combined descriptor for the *p*-th patch is given by:

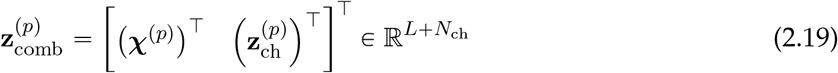

Stacking these combined descriptors from all patches row-wise yields the data matrix:

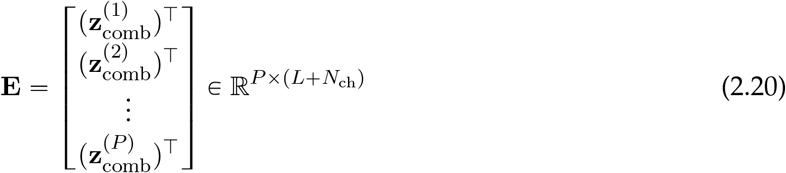

To reduce the dimensionality of the combined descriptor space while retaining most of the relevant information, we apply PCA to **E**. Let

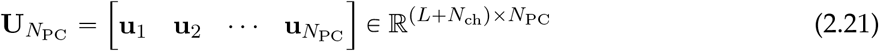

denote the loading matrix whose columns are the principal directions [59] associated with the leading modes of variation in **E**. The number of retained principal components, *N*_PC_, is chosen such that they explain over 95% of the cumulative variance. The corresponding PCA score vector for the *p*-th patch is then obtained by projecting the combined descriptor 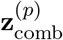 onto the loading matrix 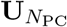, yielding

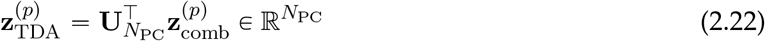

Here, 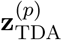 provides a reduced dimensional representation of the combined patch-wise topological and chemical descriptors. This is subsequently used to train supervised learning models for prediction of PPI sites, as discussed next.

### 2.4 Supervised Learning for Interface Prediction

The PCA-reduced descriptor for the *p*-th patch, i.e., 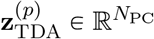, serves as the final input feature vector to the supervised learning model (NN). Thus, 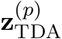 provides a compact joint representation of the local patch topology and its surrounding chemical environment.

Let 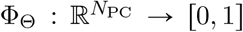, 1s denote the NN model parameterized by trainable weights and biases, collectively referred to as Θ. For the *p*-th patch, the NN predicted interface propensity is given by:

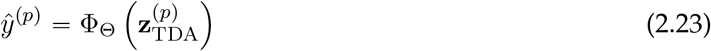

where *ŷ*^(*p*)^ ∈ [0, 1] represents the propensity (score) that the patch center belongs to an interface region. In this work, Φ_Θ_ is implemented as a feedforward multilayer NN consisting of five hidden layers with rectified linear unit (ReLU) activation functions, followed by a final sigmoid output layer to produce interface propensities, and approximately 1.8 × 10^3^ trainable parameters.

Given the true binary label *y*^(*p*)^ ∈ {0, 1} associated with the patch center, the NN parameters Θ are learned by minimizing the binary cross-entropy (BCE) loss [60, 61] over the training set, defined as:

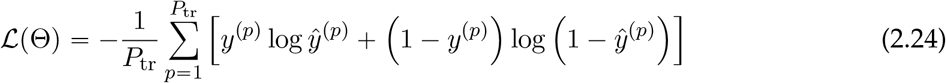

where *P*_tr_ ∈ ℤ_+_ denotes the total number of training patches. To account for the imbalance between interface and non-interface surface points, each training epoch is constructed using balanced sampling, in which an equal number of interface and non-interface patches are selected from the training proteins (similar to the MaSIF-site approach [4]). The NN model is trained using the Adam optimizer [62] following the protein-level training/testing protocol described in Section S.1 of *Supporting Information*. All surface patches extracted from the 3003 training proteins are used for model training, while all surface patches extracted from the 359 test proteins are considered for testing. No formal hyperparameter optimization is performed; the architecture is selected to provide a lightweight supervised classifier for the PCA-reduced descriptor space, while using standard training choices consistent with the MaSIF-site benchmark [4] wherever applicable, including patch-wise interface labels, binary cross-entropy loss, and Adam optimization.

To evaluate the classification performance of the proposed framework, we use the area under the receiver operating characteristic curve (AUC) as the primary performance metric [63, 64]. AUC quantifies the ability of the model to distinguish interface patches, *y*^(*p*)^ = 1, from non-interface patches, *y*^(*p*)^ = 0, over all possible classification thresholds, thereby providing a threshold-independent measure of ranking performance. Specifically, for a threshold *τ* ∈ [0, 1], the true-positive rate (TPR) and false-positive rate (FPR) are defined as:

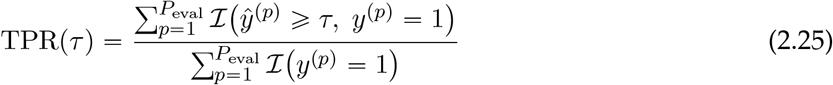

and

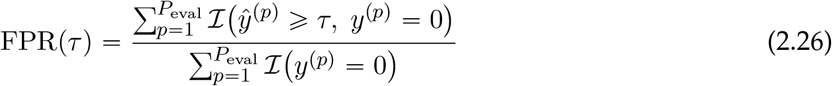

respectively, where *P*_eval_ ∈ ℤ_+_ denotes the number of patches used in evaluation and *I*(·) is the indicator function. The receiver operating characteristic (ROC) curve [65] is then obtained by tracing TPR(*τ*) against FPR(*τ*) over all thresholds *τ*, and the corresponding AUC is given by:

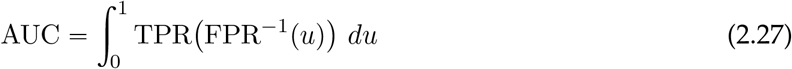

In this work, the AUC is computed as a binary score using the predicted interface propensities and the corresponding interface labels at the patch level. More specifically, the interface score is taken as the NN outputs, and the AUC is evaluated over the selected interface and non-interface patch samples used during training and testing. Therefore, higher AUC values indicate higher accuracy of the learned classifier in separating interface from non-interface patches, and this metric is used in this study for comparison with representative deep geometric learning baselines, for example, MaSIF [4]. The codes and data necessary to reproduce and implement the proposed TDA-NN framework on the PDB dataset are available at https://github.com/zavalab/ML/tree/master/TDA4Protein.

## 3 Results and Discussion

We first perform a preliminary descriptor-screening analysis on a subset of 10 randomly selected proteins from the PDB dataset. Using the moving-patch construction described in Section 2.1, we extract a total of 32,536 localized surface patches and evaluate representative TDA descriptors introduced in Section 2.2. In particular, this preliminary screening uses only the TDA descriptors computed from the patch point clouds and does not include chemical descriptors. For consistency across patches of different sizes, the EC functions are normalized by the number of points contained within the corresponding patch. Figure 4 summarizes the alpha-complex EC descriptors and their corresponding two-component PCA projection as a representative example. Additional descriptor results obtained from the rips, cubical, and PL representations are provided in Section S.3 of the *Supporting Information*.

**Figure 4.**
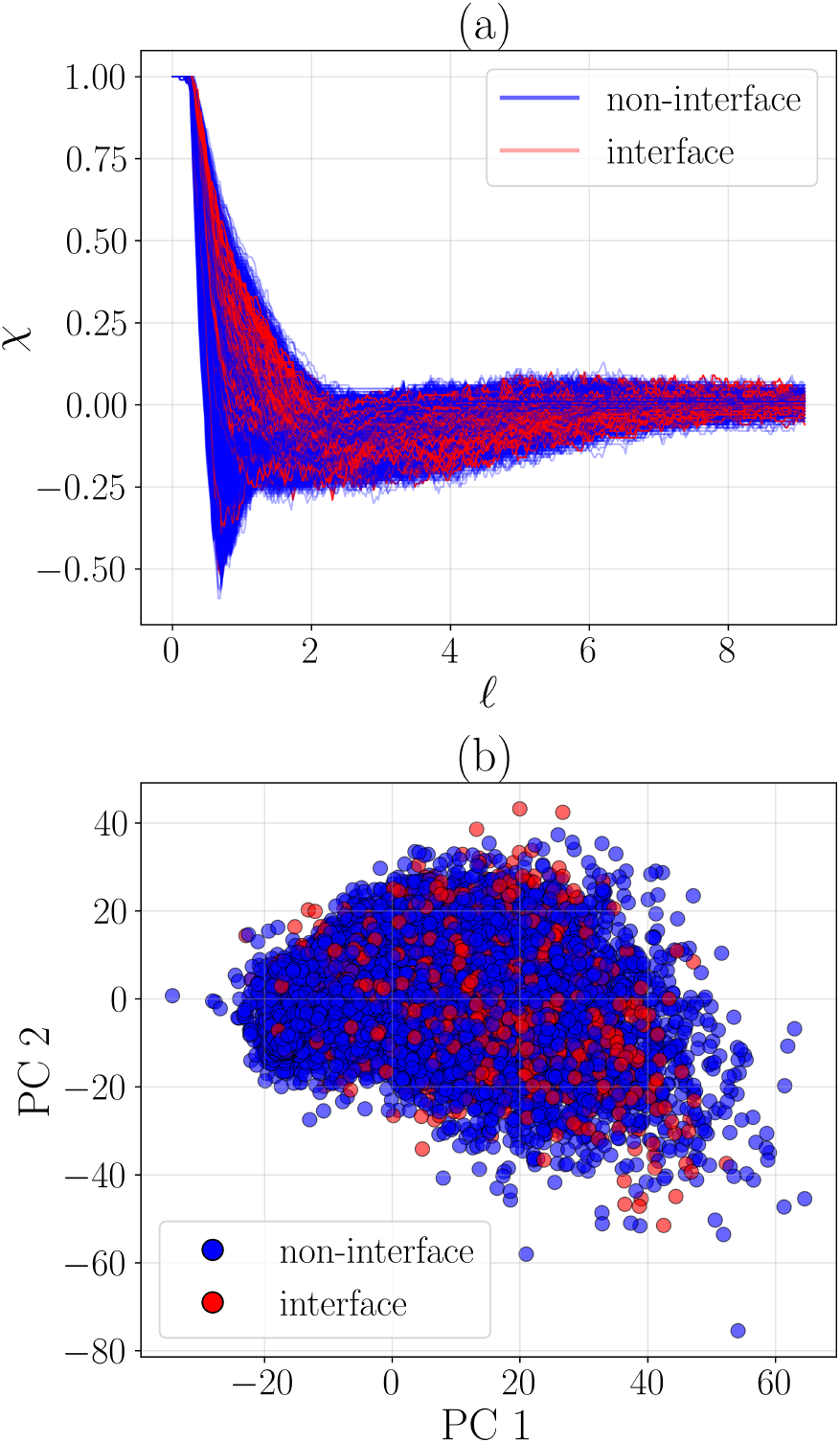
Representative EC descriptors obtained from alpha complex construction for localized protein surface patches. (a) Normalized EC functions computed from alpha complexes. (b) Two-component PCA projection of the corresponding EC descriptors, with interface patches shown in red and non-interface patches shown in blue. The PCA projection does not show a clear clustering or separation between interface and non-interface patches, suggesting that TDA descriptors might require to be coupled with supervised learning and/or augmentation with chemical descriptors to effectively discriminate PPI-prone surface regions.

Although the normalized EC functions p*χ*q obtained from alpha complex construction exhibit nontrivial variability across surface patches, the corresponding two-component PCA projection does not show a clear separation between interface and non-interface patches. Similar behavior is observed for the other TDA descriptor classes considered in this work as shown in the *Supporting Information*. These results indicate that low-dimensional unsupervised projections of TDA descriptors alone are insufficient to distinctly cluster interface and non-interface regions from localized 3D patch point clouds, thus motivating the use of supervised learning and chemically augmented descriptors for interface prediction.

To compare the candidate TDA descriptors more systematically, we next evaluate the trade-off between computational expense and predictive performance using a standard support vector machine (SVM) classifier. This preliminary analysis is intended as a warm-start study for selecting the most appropriate topological descriptor for protein surface patches prior to training the proposed topological data analysis - neural network (TDA-NN) framework on the full dataset. For each type of TDA descriptor, we retain the number of principal components required to explain more than 95% of the total variance and use these reduced descriptors as inputs to the SVM classifier. The patch radius is fixed at 9 Å for all cases. Note that this radius is the Euclidean radius of the sphere used to collect neighboring surface-mesh vertices around each patch center, rather than a geodesic radius along the molecular surface as used in MaSIF-based methods [4]. Table 1 reports the resulting per-patch computational cost, total CPU time over all 32,536 patches, and the corresponding SVM classification accuracy. For reference, the SVM accuracy obtained on the same dataset using the MaSIF geometric descriptor (shape-index and distance-dependent curvature) values [4] as input features is approximately 52.6%.

**Table 1:** Preliminary screening of candidate TDA descriptors on 32,536 localized surface patches extracted from 10 randomly selected proteins. All computations were performed on a local CPU using parallel execution with 12 concurrent workers.

| Descriptor | Time / patch (s) | Total CPU time (hr) | SVM accuracy (%) |
| --- | --- | --- | --- |
| Vietoris-Rips complex | 1.32 | 11.9 | 58.3 |
| Alpha complex | 0.02 | 0.20 | 61.1 |
| Cubical complex | 0.14 | 1.26 | 59.5 |
| Persistence landscapes | 0.03 | 0.28 | 56.8 |
| MaSIF [4] geometric descriptors | 0.43 | 3.9 | 52.6 |

Among the candidate descriptors, the alpha complex representation provides the best overall trade-off between accuracy and computational expense, achieving the highest SVM accuracy (61.1%) while remaining consistently faster than the other candidate descriptors. The comparison between MaSIF geometric descriptor baseline and the TDA descriptors in terms of classification accuracy indicates that TDA contains richer discriminatory information under the same classifier hyperparameters. Furthermore, the computation of TDA descriptors from alpha and cubical complexes, together with PL descriptors, is significantly faster than the corresponding MaSIF preprocessing workflow. Based on this preliminary analysis, we select the TDA descriptors generated from the alpha complex for all subsequent analyses.

We next scale the proposed TDA-NN framework, that combines both topological and chemical descriptors as discussed in Sections 2.3 and 2.4, to the full dataset of 3,362 proteins. To remain consistent with MaSIF-site as the benchmark framework considered in this study, we cap the maximum number of points per patch at 100. For analyzing the structure of the resulting descriptor space, Figure 5 shows a two-component PCA projection of the combined topological and chemical descriptors for all 9 Å radius patches, together with the cumulative explained variance as a function of the number of retained principal components.

**Figure 5.**
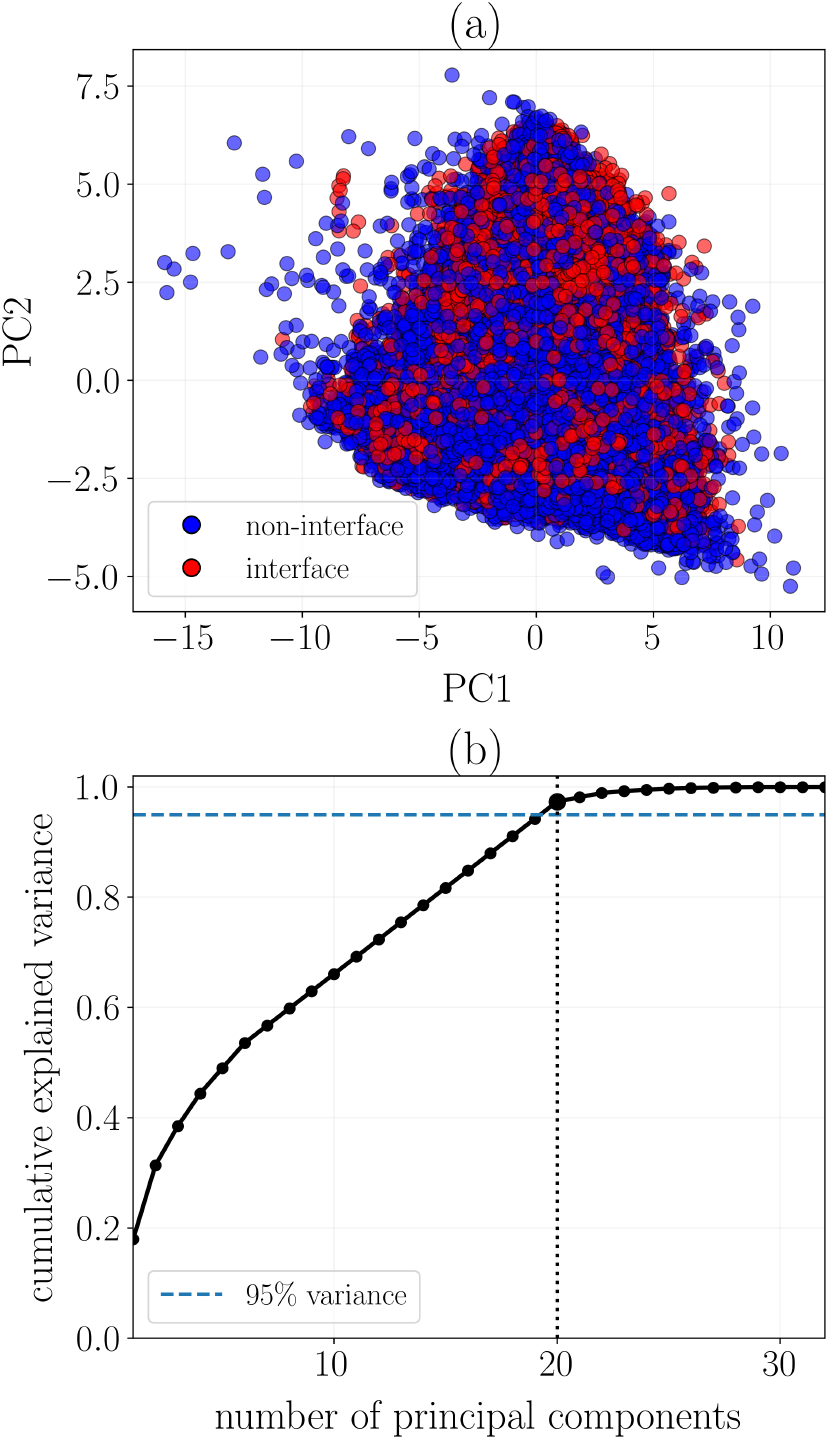
PCA of the combined descriptor space for localized protein surface patches using a 9 Å patch radius. (a) Two-component PCA projection of the combined descriptor vectors, with interface and non-interface patches shown in red and blue, respectively. The overlap between the two classes indicates that the first two principal components alone are insufficient to distinctly separate interface and non-interface patches. (b) Cumulative explained variance as a function of the number of retained principal components. The dashed horizontal line indicates the 95% cumulative-variance threshold, and the dotted vertical line indicates that 20 principal components are retained for NN training in the 9 Å case.

Although the first two principal components provide a useful low-dimensional visualization of the descriptor space, they do not distinctly separate interface and non-interface patches. This is expected because the first two principal components account for only a limited fraction (around 30%) of the total variance, while a larger number of principal components is required to retain the dominant variability in the combined descriptor space. Accordingly, for NN training, we retain the number of principal components required to explain more than 95% of the cumulative variance. One additional advantage of the proposed TDA-NN framework is its improved *scalability*. Unlike conventional geometric deep learning frameworks such as MaSIF [4] or dMaSIF [18], the TDA workflow is readily amenable to larger patch radii without a substantial increase in overall computational burden. Therefore, we consider two patch-radii values for the proposed TDA-NN framework, i.e., 9 Å and 12 Å, and compare the resulting interface prediction scores against MaSIF-site. Additional details on the implementation and usage of MaSIF-site in this work are provided in Section S.4 of the *Supporting Information* document.

The TDA-NN model is trained in a supervised manner using patch-wise interface/non-interface labels and PCA-reduced TDA and chemical descriptors as inputs, as described in Section 2.4. Figures 6 and 7 show representative qualitative comparisons between the proposed TDA-NN framework and MaSIF-site for two randomly selected proteins from the testing set. In both cases, the predicted PPI interface propensities from the TDA-NN model recover the broad spatial locations of the true interface regions and produce surface-level predictions that are qualitatively comparable to those from MaSIF-site, thus preserving the dominant PPI regions while requiring significantly reduced preprocessing and training time.

**Figure 6.**
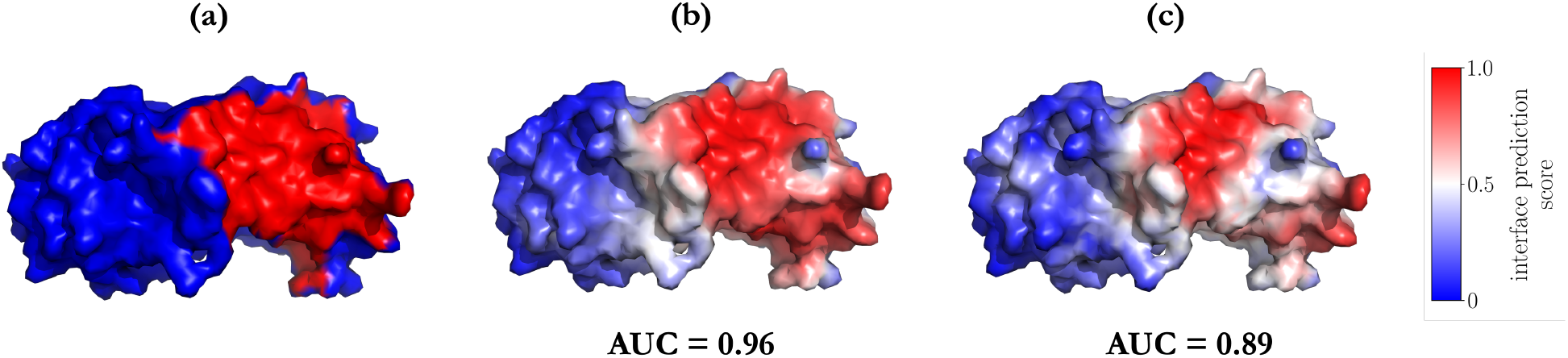
Qualitative comparison between the TDA-NN framework and MaSIF-site for a randomly selected protein (ID: `1SUW`, chain D) from the testing set using a patch radius of 9 Å. (a) Ground-truth molecular surface representation of the protein under consideration. (b) Surface-level PPI interface propensity predicted by MaSIF-site. (c) Surface-level PPI interface propensity predicted by the TDA-NN framework.

**Figure 7.**
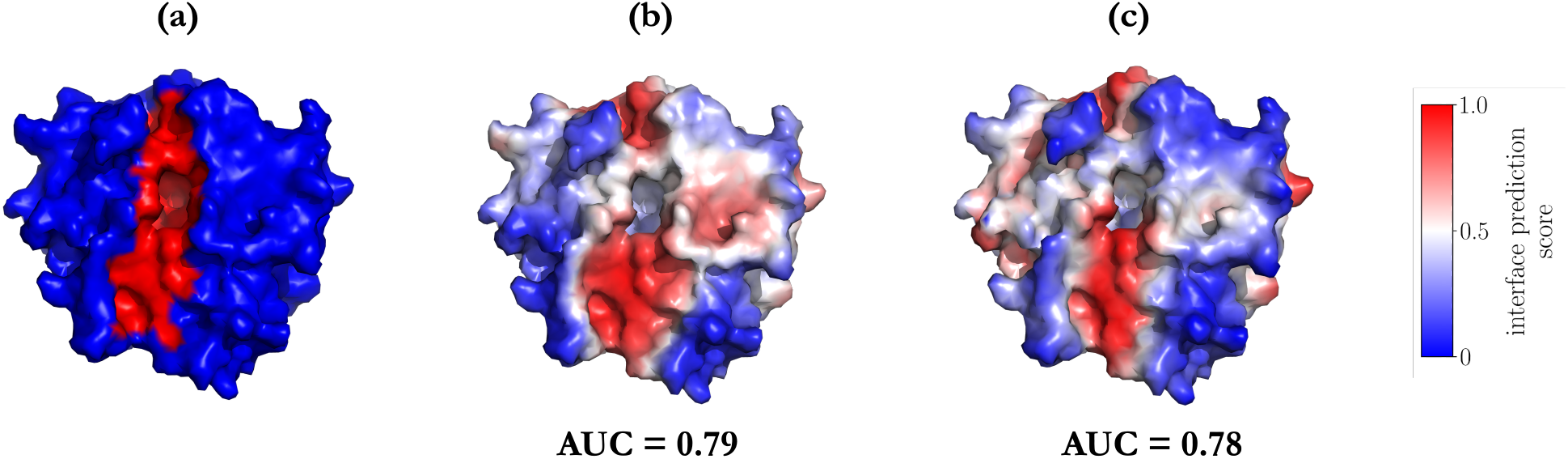
Qualitative comparison between the proposed TDA-NN framework and MaSIF-site for a randomly selected protein (ID: `2QXV`, chain A) from the testing set using a patch radius of 9 Å. (a) Ground-truth molecular surface representation of the protein under consideration. (b) Surface-level PPI interface propensity predicted by MaSIF-site. (c) Surface-level PPI interface propensity predicted by the TDA-NN framework.

To quantify these trade-offs at scale, Table 2 compares the proposed TDA-NN framework against MaSIF-site in terms of preprocessing time, total training time, and mean train/test AUC. The reduction in computational cost is accompanied by a substantially smaller TDA-NN model containing 1, 799 trainable parameters, compared with 9, 267 trainable parameters for the corresponding 9 Å MaSIF-site model. In this work, preprocessing time includes all computations performed prior to model training, such as construction of local patches, extraction of topological descriptors, and PCA of the combined topological-chemical descriptor space. To maintain a consistent basis for comparison, all three frameworks are trained for 50 epochs. In practice, however, the TDA-NN training curves saturate much earlier, with no substantial improvement after approximately 14-16 epochs for both 9 Å and 12 Å cases; similarly, MaSIF-site shows no meaningful improvement after approximately 29 epochs. The per-protein ROC AUC curves during training are provided in Figure S4 under Section S.5 in *Supporting Information*. Here, per-protein ROC AUC denotes the ROC AUC computed separately for each protein from its patch-level interface labels and predicted interface propensities. The corresponding total training times to saturation are also included in Table 2.

**Table 2:** Comparison of predictive performance and computational expense between MaSIF-site and the proposed TDA-NN framework on the full dataset.

| Framework | Patch radius (Å) | Pre-processing time | Total training time | Training time to saturation | Train AUC (mean) | Test AUC (mean) |
| --- | --- | --- | --- | --- | --- | --- |
| MaSIF-site | 9 | 27 s/protein<br>(~25.2 hr total) | 6 hr<br>(~7.2 min/epoch) | 3.5 hr<br>(~29 epochs) | 0.85 | 0.84 |
| TDA-NN | 9 | 5 s/protein<br>(~4.7 hr total) | 1 hr<br>(~1.2 min/epoch) | 16.8 min<br>(~14 epochs) | 0.77 | 0.76 |
| TDA-NN | 12 | 8 s/protein<br>(~7.5 hr total) | 1.3 hr<br>(~1.6 min/epoch) | 25.6 min<br>(~16 epochs) | 0.78 | 0.77 |

To further investigate the relative contribution of the descriptor classes, we performed an ablation study in which the NN model is trained using only chemical descriptors or only TDA descriptors, while keeping the same train-test split and training protocol. Table 3 summarizes the resulting performance for both 9 Å and 12 Å patch radii. The corresponding training ROC AUC curves from the ablation study are provided in Figure S5 under Section S.5 in *Supporting Information*. The results show that both chemical and TDA descriptors contribute meaningful information for PPI interface prediction. The TDA-only models provide competitive AUC values relative to the chemical-only models, supporting the ability of the proposed topological descriptors to capture interface-relevant geometric structure from localized surface patches. Although adding chemical descriptors to the TDA feature space produced little or no improvement in mean training AUC relative to the TDA-only feature set, the combined descriptor feature space led to slightly higher mean test AUC than either the chemical-only or TDA-only feature set in the ablation study. This result suggests that chemical descriptors may contribute more to model generalization than to fitting the training data. This observation is also consistent with the ablation study in the original MaSIF work, which showed that chemical surface features were more important than geometric features for characterizing PPI patterns [4].

**Table 3:** Ablation study evaluating the relative contribution of chemical and TDA descriptors for interface prediction using the NN model.

| Feature set | Patch radius<br>(Å) | Train AUC<br>(mean) | Test AUC<br>(mean) |
| --- | --- | --- | --- |
| Chemical only | 9 | 0.76 | 0.74 |
| TDA only | 9 | 0.78 | 0.73 |
| Chemical only | 12 | 0.77 | 0.75 |
| TDA only | 12 | 0.78 | 0.73 |

The quantitative results confirm the same trend observed qualitatively in Figures 6 and 7. MaSIF-site achieves the highest predictive accuracy on both the training and testing sets, but at significantly higher preprocessing and training times. In contrast, the proposed TDA-NN framework yields only a modest decrease in train/test AUC while providing a substantial improvement in scalability and computational efficiency. For the 9 Å patch radius, the TDA-NN model reduces the average preprocessing time from approximately 27 s/protein to 5 s/protein and the total training time from 6 h to 1 h, while maintaining a test AUC of 0.76. Increasing the patch radius to 12 Å produces a slight improvement in predictive performance, with a test AUC of 0.77, at the expense of a moderate increase in preprocessing and training times. These results demonstrate that the proposed TDA-based representation can accommodate larger local neighborhoods with manageable additional cost, which can be considered as an attractive property for large-scale interface analysis.

Overall, these results pose the proposed TDA-NN framework as a computationally efficient alternative to benchmark geometric deep learning pipelines for PPI interface prediction. Compared to MaSIF, the TDA-NN approach achieves reasonable predictive accuracy at substantially lower preprocessing and training costs. dMaSIF addresses related computational bottlenecks of MaSIF through a distinct acceleration strategy, in which molecular-surface points, features, and local coordinate systems are computed on the fly from atomic point clouds [18]. In contrast, the present work focuses on a different simplification strategy: localized molecular-surface patches are compressed into compact topological descriptors prior to training, avoiding the need for learned quasi-geodesic convolutions on surface patches. A direct numerical comparison with dMaSIF is therefore outside the scope of the present study, as it would require implementing and benchmarking both the original dMaSIF workflow and a corresponding TDA-augmented dMaSIF variant under a consistent surface representation, feature set, and training protocol. In this sense, the proposed method complements frameworks such as MaSIF and dMaSIF by emphasizing scalability and fast computation, while still maintaining competitive accuracy for large-scale structure-based interface prediction.

## 4 Conclusions and Future Work

In this work, we introduced a scalable topological data analysis (TDA) framework for protein-protein interface prediction from localized protein surface patches. Starting from protein molecular surface point clouds, the proposed workflow constructs local point cloud patches, evaluates multi-scale topological descriptors, and combines these descriptors with chemical information for supervised interface prediction using multilayered neural networks (NN). On the full dataset, the proposed TDA-NN framework achieved competitive train/test AUC values while requiring substantially lower preprocessing and training times than benchmark geometric deep learning approaches such as MaSIF. These results indicate that TDA provides a scalable and computationally efficient alternative to conventional geometric deep learning frameworks for large-scale protein interface prediction. Although MaSIF achieves higher predictive accuracy, the proposed TDA-NN framework demonstrates that competitive interface prediction can be achieved with substantially faster preprocessing and training, while naturally accommodating larger patch radii and large-scale datasets.

Future work will focus on extending the proposed framework in several directions. First, weighted persistent homology formulations can be incorporated so that chemical information is used directly during the computation of TDA descriptors through weighted filtrations, rather than being appended afterward in the input feature space [43]. This readily enables the integration of topology and physicochemical heterogeneity within a unified descriptor construction. Second, the same computational framework can be extended beyond PPI prediction to related biomolecular problems, such as protein-ligand binding site prediction and other structure-based molecular recognition tasks [66, 41]. These directions will further clarify the potential of TDA as a scalable and computationally efficient paradigm for biomolecular interface modeling.

## Data Availability Statement

The protein structures and PPI datasets analyzed in this study were obtained from publicly available sources, as described in the manuscript. The processed data and source code required to reproduce the preprocessing, topological descriptor computation, model training, and evaluation procedures are publicly available at https://github.com/zavalab/ML/tree/master/TDA4Protein.

## Supporting Information

Description of the PPI datasets; molecular-surface generation, chemical-feature calculation, and interface-label assignment procedures; additional visualizations of topological descriptors; implementation details for the MaSIF-site benchmark; and additional TDA-NN results, including training curves and feature-ablation analyses (PDF).

## Acknowledgements

This work was funded though the National Science Foundation under award number 2346683.

