## Supporting Information for "Scalable Extraction of Information on Protein-Protein Interactions using Topological Data Analysis"

The following items are provided in the supporting information document for clarity.

- **S.1**: Description of Protein-Protein Interaction Datasets
- **S.2**: Data Preprocessing Steps
- **S.3**: Additional Visualizations of Topological Descriptors
- **S.4**: Additional Details on MaSIF-site Implementation
- **S.5**: Additional Results from the TDA-NN Framework

### S.1 Description of PPI Datasets

Protein-protein interaction (PPI) pairs were obtained from the MaSIF-site dataset [1], which was taken from the PRISM nonredundant protein set [2], the ZDock benchmark [3], PDBBind [4], and SabDab [5]. In total, 3362 proteins were used in this study, comprising 3003 proteins for training and 359 proteins for testing. Since interface prediction is performed for individual surface patches rather than whole proteins, this protein-level split implies that all surface patches extracted from a training protein are used only for training, while all surface patches extracted from a test protein are used only for testing.

The training and test sets followed the sequence- and structure-based split defined in the original MaSIF-site [1] study. Specifically, this split was designed to reduce overlap between the training and testing proteins by excluding closely related proteins and structurally similar interaction sites from appearing across both sets. Thus, the test set provides a more stringent assessment of interface prediction performance on proteins and binding-site geometries that are not trivially represented in the training set. Additional details regarding dataset construction and splitting criteria are provided in [1].

### S.2 Data Preprocessing Steps

#### S.2.1 Computation of discretized molecular dot surfaces

For consistency with the original MaSIF-site study, molecular dot surfaces were computed using MSMS [6] with the same parameters: surface density = $3.0$ and water probe radius = $1.5$ Å. Although subsequent preprocessing into a discretized triangulated surface mesh was not required for the computation of topological descriptors, triangulated molecular surfaces were generated to visualize predicted interfaces on protein surface representations (as represented in Figures 5a and 6a in the manuscript). Following the original MaSIF-site workflow, the irregular meshes produced by MSMS were further regularized using PyMesh [7] (v0.3.1) at a resolution of $1.0$ Å. As a result, each protein molecular surface was represented both as a point cloud of molecular surface dots and as a discretized triangulated mesh for visualization and downstream analysis.

#### S.2.2 Chemical surface feature calculations

Each molecular surface vertex was assigned three chemical features: hydropathy index [8], Poisson-Boltzmann electrostatic potential [9], and hydrogen bond potential [10, 11]. These features and their calculation methods match those used in MaSIF-site [1] and are summarized as follows.

- *Hydropathy index*: Each molecular surface vertex was assigned a hydropathy value based on the Kyte-Doolittle scale [8], according to the amino acid identity of the nearest atom. The original scale, ranging from $-4.5$ for the most hydrophilic residues to $+4.5$ for the most hydrophobic residues, was normalized to the range $[-1, 1]$.

- *Poisson-Boltzmann electrostatic potential*: PDB2PQR [12] was used to prepare protein structures for electrostatic calculations, and Poisson-Boltzmann electrostatics were computed using APBS [13] (v.1.5). The electrostatic potential at each vertex of the triangulated molecular surface was assigned using Multivalue, a utility within the APBS suite [13]. Potential values greater than $+30$ or less than $-30$ were capped at these limits, and the resulting values were normalized to the range $[-1, 1]$.

- *Hydrogen bond potential*: Free-electron positions and potential hydrogen bond donors on the molecular surface were computed using the hydrogen bond potential model described in [10]. Surface vertices whose nearest atom was a polar hydrogen, nitrogen, or oxygen were identified as potential hydrogen-bond donor or acceptor sites. Each vertex was then assigned a value from a Gaussian-shaped potential function based on the orientation of the relevant heavy atoms. The resulting values range from $-1$, corresponding to an optimal position for a hydrogen bond acceptor, to $+1$, corresponding to an optimal position for a hydrogen bond donor.

##### S.2.3 Definition of PPI interface points on protein surface dots

Ground-truth labels for protein surface dots were defined by comparing the molecular surface of each isolated monomer with the molecular surface of its corresponding bound complex obtained from the RCSB Protein Data Bank [14]. Because interface regions become buried and are no longer solvent-exposed upon complex formation, monomer surface dots that did not overlap with the surface points of the bound complex were assigned positive interface labels. All remaining surface dots were assigned negative non-interface labels.

#### S.3 Additional Visualizations of Topological Descriptors

This section includes additional TDA descriptor results obtained from the rips, cubical, and PL representations of the 32,536 localized surface patches (protein point clouds). For consistency across patches of different sizes, the EC functions are normalized by the number of points contained within the corresponding patch.

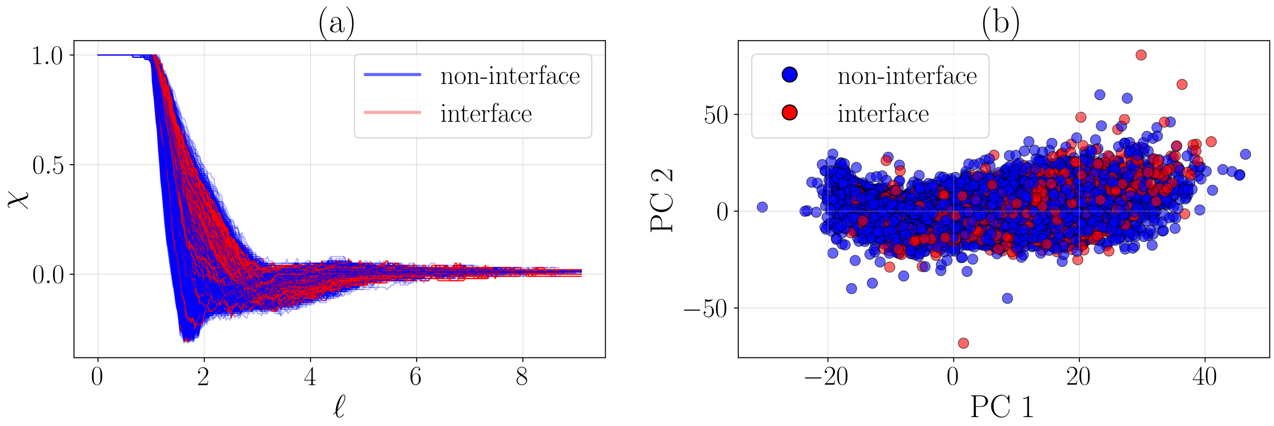


Figure S1: Representative EC descriptors obtained from rips complex construction for localized protein surface patches. (a) Normalized EC functions computed from rips complexes. (b) Two-component PCA projection of the corresponding EC descriptors, with interface patches shown in red and non-interface patches shown in blue.

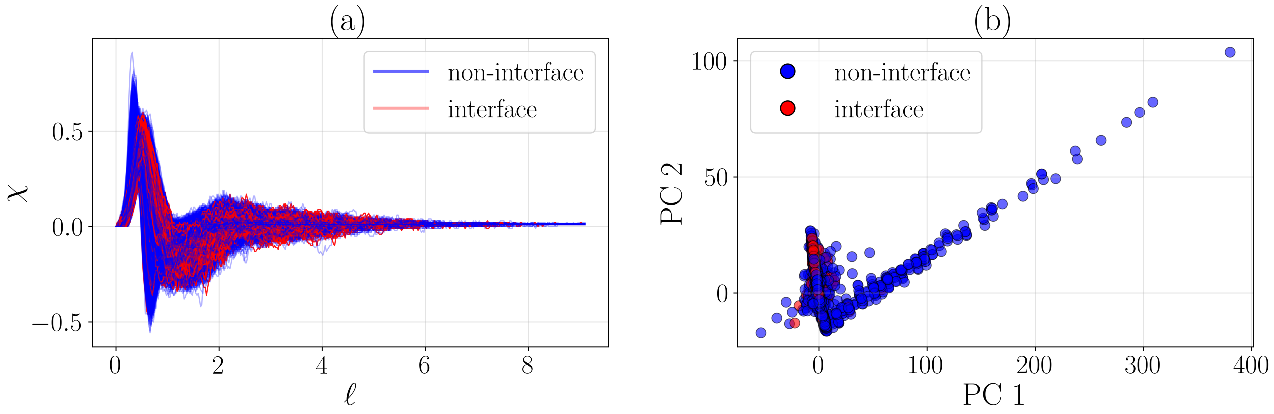


Figure S2: Representative EC descriptors obtained from cubical complex construction for localized protein surface patches. (a) Normalized EC functions computed from cubical complexes. (b) Two-component PCA projection of the corresponding EC descriptors, with interface patches shown in red and non-interface patches shown in blue. The cubical-complex PCA projection exhibits a more structured low-dimensional pattern than the rips- and alpha-complex projections. This likely arises from the grid-based distance-transform filtration, which imposes a smoother and more ordered evolution of sublevel sets across patches. However, interface and non-interface patches still remain substantially overlapped.

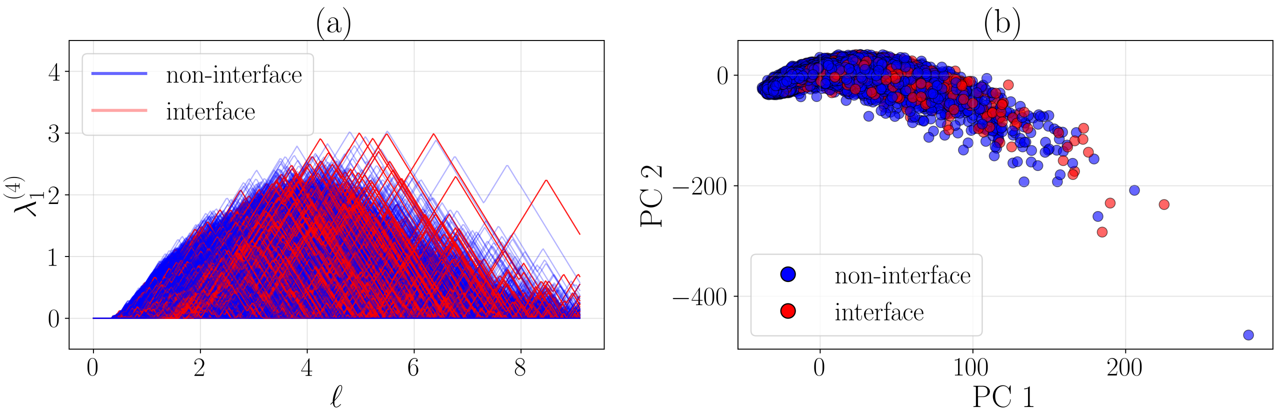


Figure S3: Representative PL descriptors obtained from localized protein surface patches. (a) Representative PL functions for homological dimension $q = 1$ and landscape level $k = 4$. (b) Two-component PCA projection of the corresponding PL descriptors constructed from all retained landscape levels $k = 1, 2, \ldots, 5$, with interface patches shown in red and non-interface patches shown in blue.

#### S.4 Additional Details on MaSIF-site Implementation

The MaSIF-site model was reimplemented in PyTorch v2.1.2 following the architecture of the original MaSIF-site model [1]. This reimplementation was developed to improve scalability, training efficiency, ease of implementation, and downstream analysis. Consistent with the original architecture, the model consisted of three convolutional layers that operated on mapped surface-patch features.

The reimplemented model achieved prediction performance comparable to that of the original MaSIF-site model. Specifically, it yielded a median per-protein ROC AUC of 0.86 across the full test set, compared with 0.87 reported for the original implementation. This comparable performance supports its use as a benchmark for comparison with our TDA descriptor-based approach. The model was trained using the same key parameters as the original study: a patch geodesic radius of 9 Å, a maximum of 100 surface vertices per patch, and 43 training epochs. Additional details regarding the model architecture and training parameters are provided in [1].

For patch-wise interface prediction, two geometric surface features used in the original MaSIF-site model were precomputed in addition to the chemical features described in Section S.2.2.

- *Shape index*: The shape index (Equation S.1) describes the local geometry around each surface point based on its curvature [15]. Its values range from $-1$, corresponding to highly concave regions, to $+1$, corresponding to highly convex regions. The shape index is defined in terms of the principal curvatures $\kappa_1$ and $\kappa_2$, where $\kappa_1 \geqslant \kappa_2$, as follows:

$$\text{Shape index} = \frac{2}{\pi} \tan^{-1}\left(\frac{\kappa_1 + \kappa_2}{\kappa_1 - \kappa_2}\right) \tag{S.1}$$

  The principal curvatures were calculated from the mean curvature $H$ and Gaussian curvature $K$ according to

$$\kappa_1 = H + \sqrt{H^2 - K}, \qquad \kappa_2 = H - \sqrt{H^2 - K} \tag{S.2}$$

Both the vertex mean curvature $H$ and vertex Gaussian curvature $K$ were computed using PyMESH [7].

- *Distance-dependent curvature*: For each vertex within an extracted surface patch, the distance-dependent curvature assigns a value in the range $[-0.7, 0.7]$ that captures the relationship between the vertex's distance from the patch center and the orientation of its surface normal relative to the center point. Further details regarding the calculation of this feature are provided in [15]. Whereas the shape index is a principal-curvature-based descriptor that characterizes the local geometry of individual vertices across the protein surface, the distance-dependent curvature characterizes geometry at the patch level using the patch center as a reference. The distance-dependent curvature has therefore been shown to provide geometric information complementary to that captured by the shape index [1, 15].

#### S.5 Additional Results from the TDA-NN Framework

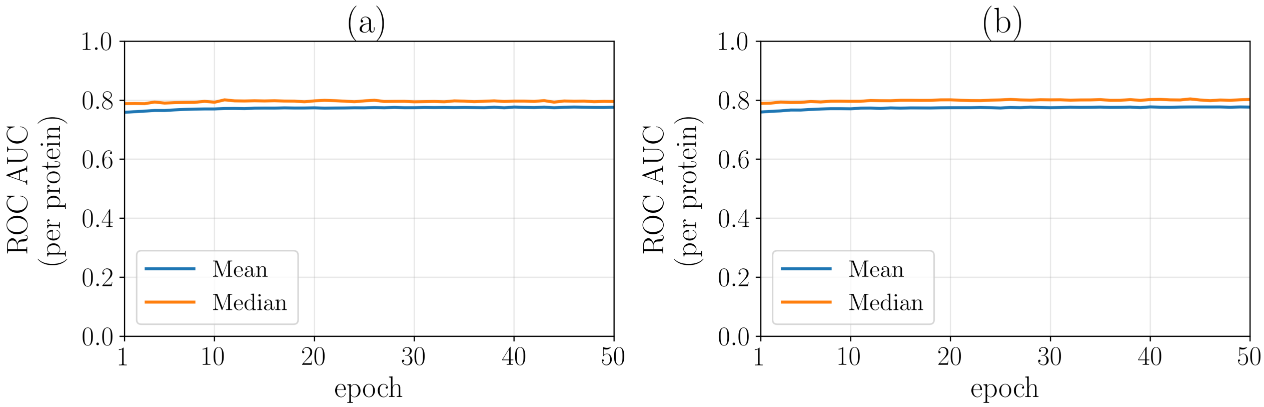


Figure S4: Training evolution of ROC AUC for the TDA-NN framework using combined topological and chemical descriptors for the (a) 9 Å and (b) 12 Å cases. At each epoch, the ROC AUC is first computed separately for each protein using the patch-level and predicted interface propensities, and the resulting mean and median values across proteins (per-protein ROC AUC) are reported. Although the NN models are trained for 50 epochs, the ROC AUC curves saturate much earlier, with no substantial improvement after approximately 14 epochs for the 9 Å case and 16 epochs for the 12 Å case.

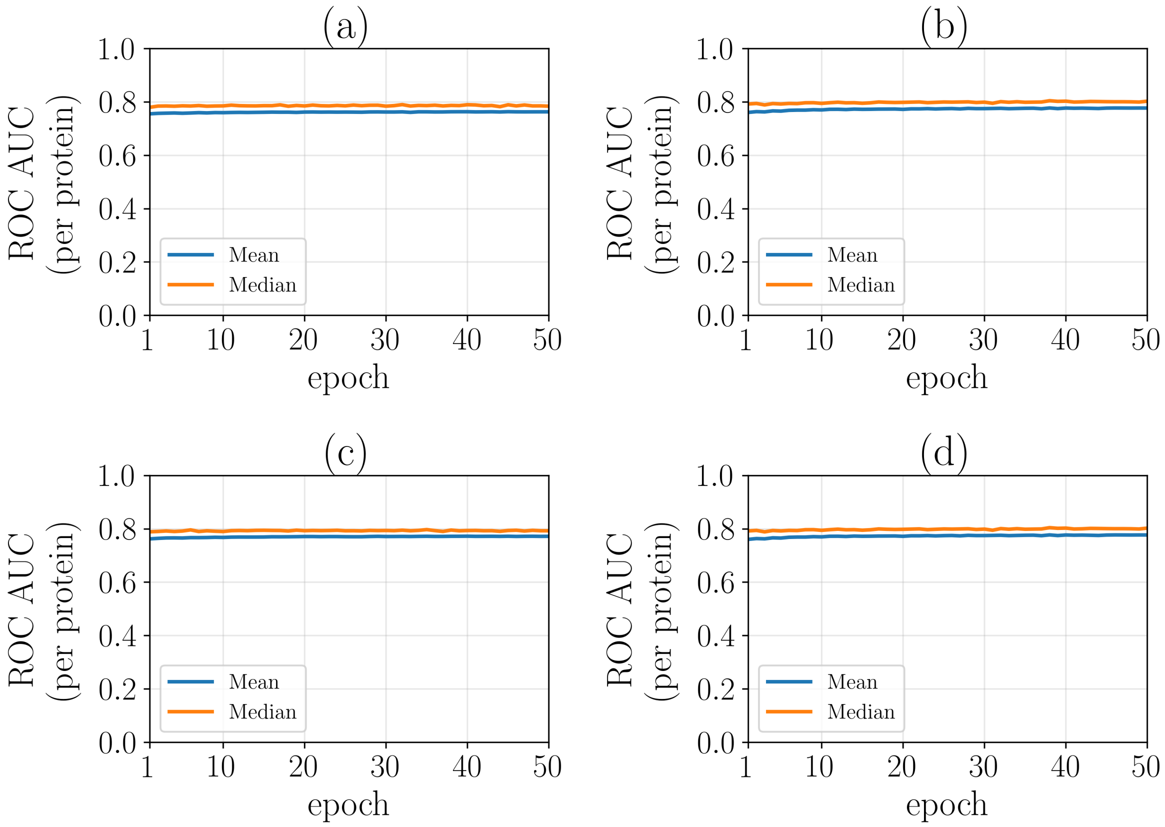


Figure S5: Training ROC AUC curves for the ablation study evaluating the relative contribution of chemical and TDA descriptors. Mean and median train ROC AUC values are shown over 50 training epochs for four cases: (a) chemical-only descriptors with a 9 Å patch radius, (b) TDA-only descriptors with a 9 Å patch radius, (c) chemical-only descriptors with a 12 Å patch radius, and (d) TDA-only descriptors with a 12 Å patch radius. The curves show that all ablated models reach stable performance within the training window.

[2] Alper Baspinar, Engin Cukuroglu, Ruth Nussinov, Ozlem Keskin, and Attila Gursoy. PRISM: a web server and repository for prediction of protein-protein interactions and modeling their 3d complexes. *Nucleic Acids Research*, 42(W1):W285–W289, 2014.

[3] Thom Vreven, Iain H. Moal, Anna Vangone, Brian G. Pierce, Panagiotis L. Kastritis, Mieczyslaw Torchala, Raphael Chaleil, Brian Jiménez-García, Paul A. Bates, Juan Fernandez-Recio, Alexandre M. J. J. Bonvin, and Zhiping Weng. Updates to the integrated protein-protein interaction benchmarks: docking benchmark version 5 and affinity benchmark version 2. *Journal of Molecular Biology*, 427(19):3031–3041, 2015.

[4] Zhihai Liu, Yan Li, Lili Han, Jie Li, Jie Liu, Zhonghua Zhao, Wen Nie, Yuchen Liu, and Renxiao Wang. PDB-wide collection of binding data: current status of the PDBbind database. *Bioinformatics*, 31(3):405–412, 2015.

[5] James Dunbar, Konrad Krawczyk, Jinwoo Leem, Terry Baker, Angelika Fuchs, Guy Georges, Jiye Shi, and Charlotte M. Deane. SAbDab: the structural antibody database. *Nucleic Acids Research*, 42(D1):D1140–D1146, 2014.

[12] Todd J. Dolinsky, Paul Czodrowski, Hui Li, Jens E. Nielsen, Jan H. Jensen, Gerhard Klebe, and Nathan A. Baker. PDB2PQR: expanding and upgrading automated preparation of biomolecular structures for molecular simulations. *Nucleic Acids Research*, 35(suppl_2):W522–W525, 2007.

[13] Nathan A. Baker, David Sept, Simpson Joseph, Michael J. Holst, and J. Andrew McCammon. Electrostatics of nanosystems: application to microtubules and the ribosome. *Proceedings of the National Academy of Sciences*, 98(18):10037–10041, 2001.

[14] Helen M Berman, John Westbrook, Zukang Feng, Gary Gilliland, Talapady N Bhat, Helge Weissig, Ilya N Shindyalov, and Philip E Bourne. The protein data bank. *Nucleic acids research*, 28(1):235–242, 2000.

[15] Shuangye Yin, Elizabeth A Proctor, Alexey A Lugovskoy, and Nikolay V Dokholyan. Fast screening of protein surfaces using geometric invariant fingerprints. *Proceedings of the National Academy of Sciences*, 106(39):16622–16626, 2009.
